# Modified 3^rd^ Generation Synchronization Modulation Electric Field (3m-SMEF) Prevents Ischemia-Reperfusion-Induced Acute Kidney Injury via Targeting Na⁺/K⁺-ATPase

**DOI:** 10.64898/2026.10.15.759393

**Authors:** Nathan Hall, Nadezhda N. Zheleznova, Carlos Esquivel, Tamara A. Wahbeh, Colby Parris, Isabella Lozonschi, Nohely Hernandez Soto, Lei Wang, Wei Chen, Ruisheng Liu

## Abstract

Acute kidney injury (AKI) caused by ischemia–reperfusion injury (IRI) remains a major clinical problem with limited therapeutic options. We previously demonstrated that the 3rd-generation Synchronization Modulation Electric Field (SMEF) preserves Na⁺/K⁺-ATPase function during ischemia and protects against kidney injury. In the present study, we substantially modified the 3rd-generation SMEF (3m-SMEF) to further improve/optimize Na⁺/K⁺-ATPase regulation and enhance the protective effects, which was tested both in vitro and in vivo. In HK-2 cells, hypoxia/reoxygenation reduced Na⁺/K⁺-ATPase expression and induced apoptotic and necroptotic signaling, whereas 3m-SMEF restored Na⁺/K⁺-ATPase expression and markedly reduced cell death. In vivo, male and female mice underwent unilateral renal ischemia with or without 3m-SMEF treatment during vascular clamping. At 24 h, 3m-SMEF markedly improved renal function, reducing plasma creatinine by ∼90–95% and restoring glomerular filtration rate to near-control levels. Application of the 3m-SMEF also normalized kidney injury markers (KIM-1 and NGAL), preserved renal ATP, and prevented the translocation and loss of Na⁺/K⁺-ATPase expression. The renoprotective effects substantially exceeded those achieved with the previous SMEF platform. These findings demonstrate that 3m-SMEF provides robust protection against ischemic AKI by preserving Na⁺/K⁺-ATPase function and tubular integrity and represents a promising non-pharmacological therapy for ischemic kidney injury.

## Introduction

Acute kidney injury (AKI) remains a major clinical problem with limited therapeutic options and high rates of morbidity and mortality ^1^. Ischemia-reperfusion injury (IRI) is one of the leading causes of AKI. One of the earliest and most critical events during IRI is the disruption of cellular energy balance, which directly impairs ion transport systems essential for tubular epithelial cell survival ^2,3^. The Na⁺/K⁺-ATPase pump is central to maintaining intracellular ion homeostasis, cell volume, and epithelial polarity. Under ischemic conditions, ATP depletion rapidly inhibits Na⁺/K⁺-ATPase activity, leading to intracellular sodium accumulation, cellular swelling, and eventual cell death ^4,5^. Loss of pump function has been strongly associated with tubular injury, increased injury markers such as kidney injury molecule-1 (KIM-1) and neutrophil gelatinase-associated lipocalin (NGAL) ^6,7^, and decline in renal function, highlighting it as a key mechanistic driver of AKI.

Targeting Na⁺/K⁺-ATPase dysfunction and restoring cellular energetics therefore represents a promising therapeutic strategy. Recently, bioelectrical modulation has emerged as an innovative approach to regulate cellular function. Electrical fields can influence membrane potential, ion transport activity, and metabolic processes ^8,9^. In our previous work, we developed a third-generation Synchronization Modulation Electric Field (SMEF) system, which demonstrated significant protection against renal IRI by maintaining Na⁺/K⁺-ATPase function and reducing the severity of kidney injury ^10^. These findings established the critical role of Na⁺/K⁺-ATPase dysfunction in the development of IRI-induced AKI. To more effectively regulate Na⁺/K⁺-ATPase activity during ischemia and thereby provide greater protection against kidney injury, we made substantial modifications to the third-generation SMEF platform (3m-SMEF). The new generation incorporates dual energy transformation functions and power injections to more efficiently generate ATP. The pumping rate and the synchronization of the pump molecules were also further optimized to minimize potential side effects on the cells. This 3m-SMEF system is intended to further reduce the dependence of Na⁺/K⁺-ATPase activity on external ATP availability and to more rapidly synchronize the pump to its target operating frequency, thereby maintaining ionic concentration gradients and the membrane potential under conditions of ATP depletion. We hypothesized that the 3m-SMEF system would more effectively preserve Na⁺/K⁺-ATPase function, better maintain cellular homeostasis, and ultimately prevent the development of AKI following IRI.

In the present study, we tested 3m-SMEF in vitro in human proximal tubular epithelial (HK-2) cells and in vivo in a mouse IRI-AKI model. We used HK-2 cells exposed to hypoxic conditions to investigate the direct cellular effects of 3m-SMEF, including modulation of hypoxia-signaling pathways such as HIF-1α ^11^. Then we used a murine model of renal IRI in both male and female C57BL/6J mice. Renal function was assessed by plasma creatinine and GFR, while tubular injury was evaluated using KIM-1 and NGAL levels. In addition, tissue ATP levels were measured to assess cellular energetics. We further examined the localization of Na⁺/K⁺-ATPase and its protein expression as a mechanistic readout of pump integrity.

## Results

### Application of the 3m-SMEF reduces hypoxia-induced necroptosis and apoptosis in HK-2 cells

We first examined the effects of the 3m-SMEF on Hypoxia/Reoxygenation (H/R)-induced injury in HK-2 cells. Under normoxic conditions, HK-2 cells showed minimal Cytox and Annexin V fluorescent staining (ab14144), indicating low basal levels of apoptosis and cytotoxicity (Fig.1A). Hypoxic exposure for 24 h markedly increased cell necroptosis and apoptosis in HK-2 cells compared with cells maintained under normal conditions, as demonstrated by increased Cytox (green) and Annexin V (red) staining (Fig.1B). Application of the 3m-SMEF during hypoxic exposure significantly attenuated hypoxia-induced cellular injury by reducing Cytox and Annexin V fluorescence intensity (Fig.1C) compared with the hypoxia group (Fig.1B), indicating decreased necroptosis and apoptosis. Quantitative analysis confirmed that hypoxia significantly increased the number of Cytox- and Annexin V-positively stained cells compared with the Normal group (about 5.1 and 5.8-fold, respectively) (Fig. 1, D and E), whereas 3m-SMEF treatment reduced these responses toward near-control levels. These data show that 3m-SMEF exerts a significant protective effect against hypoxia-induced injury in HK-2 cells.

**Figure 1.**
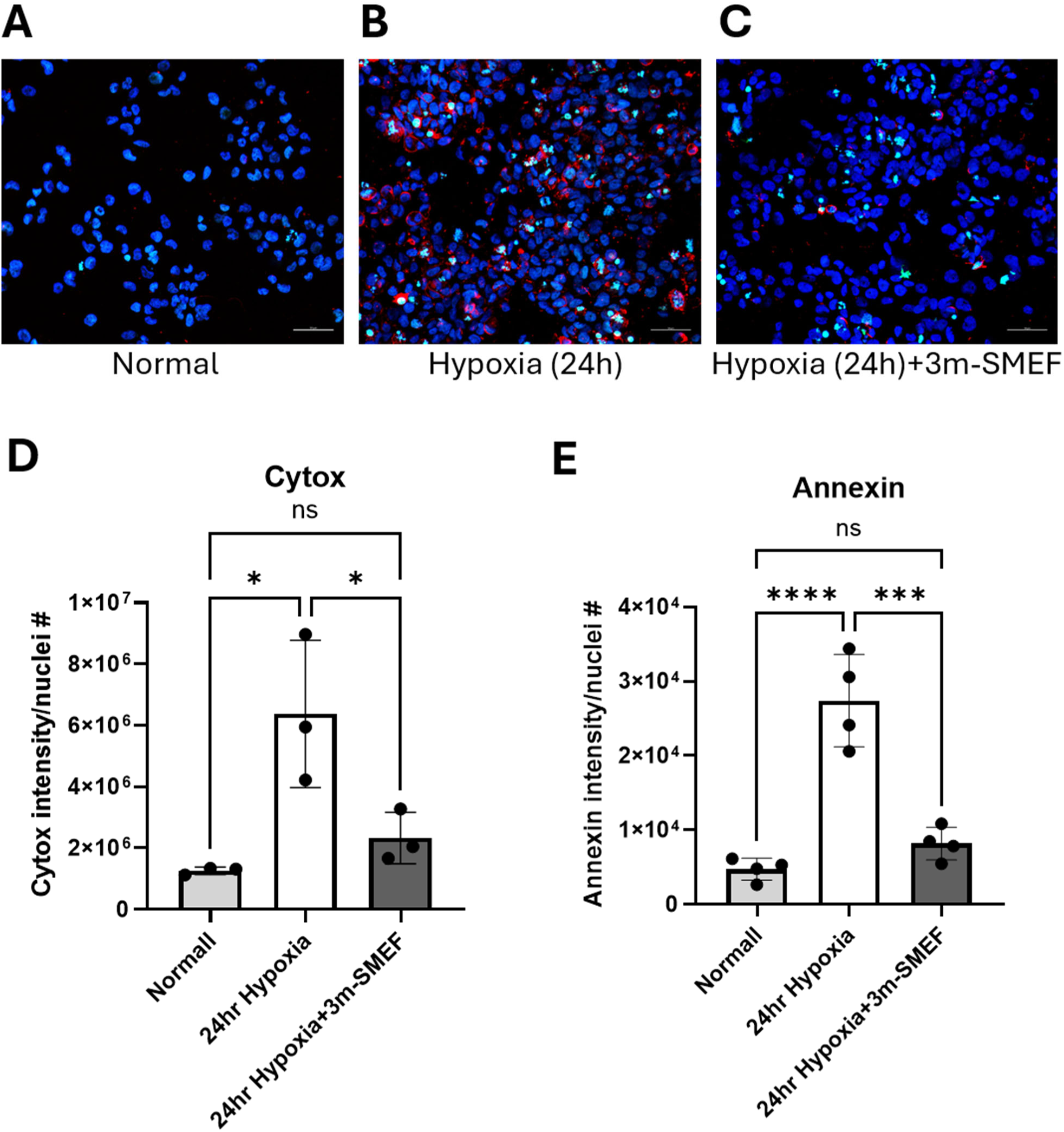
The modified third-generation SMEF (3m-SMEF) attenuates hypoxia-induced cellular injury in HK-2 cells. Representative immunofluorescence images of HK-2 cells stained with Cyto× Green (green, marker of necrotic/necroptotic cell death) and Annexin V (red, marker of apoptosis) under normal conditions *(A),* after 24 h of hypoxia *(B),* and after 24 h of hypoxia with application of 3m-SMEF *(C).* Nuclei were counterstained with DAPI (blue). Scale bar = 50 pm. Quantitative analysis of Cytox Green-positive cells *(D)* and Annexin V-positive cells *(E).* Data are presented as mean ± SEM. N=6-7. Data were analyzed using one-way ANOVA. * P < 0.05, *** P < 0.001, ****P< 0.0001.

### Application of the 3m-SMEF restores hypoxia-induced reduction of Na⁺/K⁺-ATPase in HK-2 cells

Hypoxia reduced Na⁺/K⁺-ATPase expression in HK-2 cells (Fig. 2). Immunofluorescence staining revealed internalization of Na⁺/K⁺-ATPase (red) following hypoxia, indicating impaired membrane localization and a reduction in the number of functional pumps on the cell membrane compared with the normoxic control (Fig. 2, A and B). Consistent with these findings, Western blot analysis demonstrated a marked reduction in Na⁺/K⁺-ATPase α1 protein expression under hypoxic conditions (Fig.2, D and E) suggesting hypoxia-induced downregulation and/or degradation of the pump. In contrast, application of the 3m-SMEF prevented Na⁺/K⁺-ATPase internalization and restored Na⁺/K⁺-ATPase α1 expression to levels exceeding those observed in normoxic/normal controls (Fig. 2, D and E). This finding suggests that synchronized electrical stimulation (3m-SMEF) not only preserves pump stability and membrane localization during hypoxic stress but may also enhance Na⁺/K⁺-ATPase abundance.

**Figure 2.**
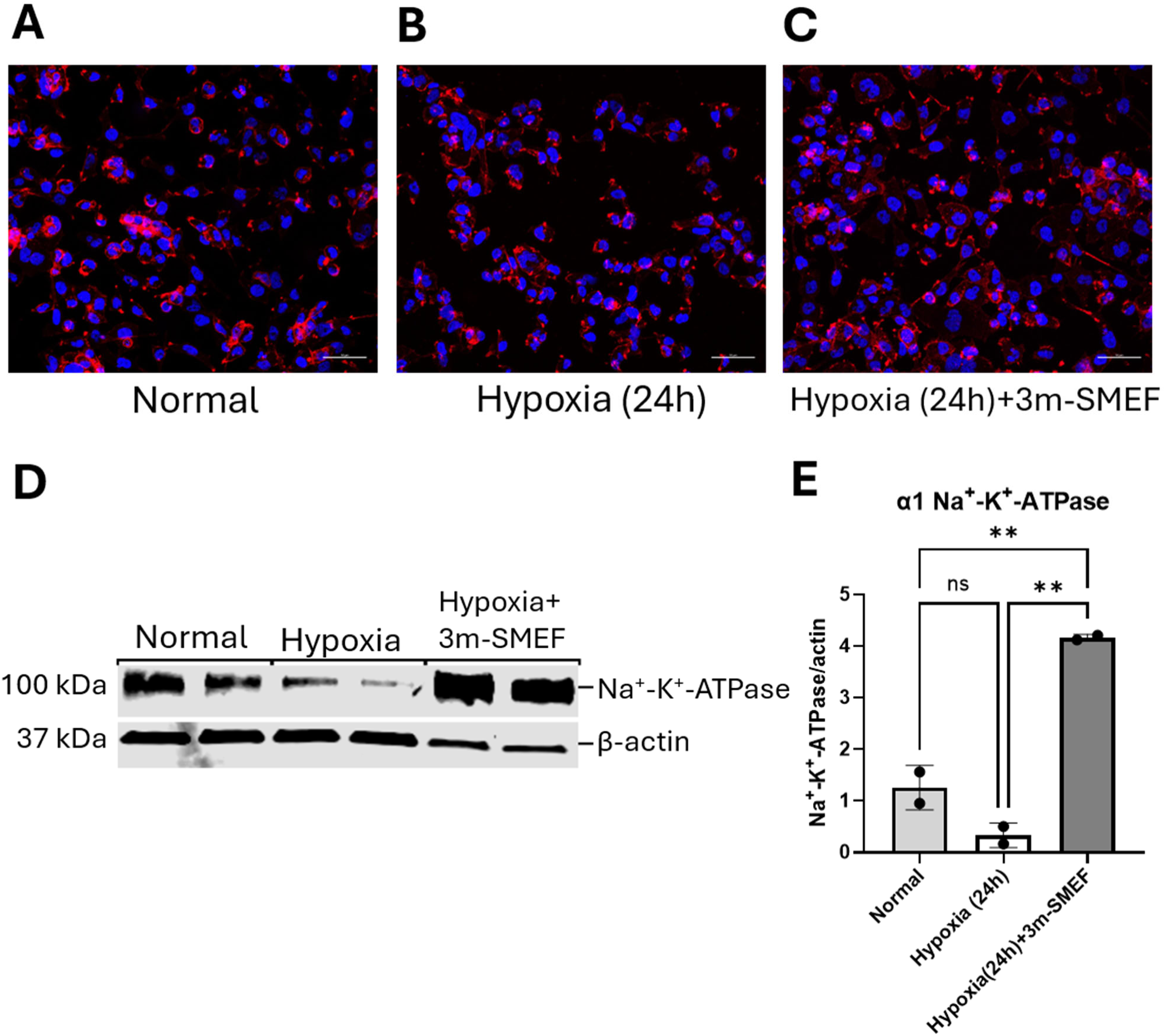
Effect of 3m-SMEF treatment on hypoxia-induced changes in expression on Na^+^-K^+^-ATPase in HK-2 cells. Representative immunofluorescence staining of HK-2 cells with Na^+^-K*-ATPase antibodies (red staining) under normal conditions *(A),* hypoxia (24h) *(B),* and hypoxia (24h) with 3m-SMEF application *(C).* Nuclei were counterstained with DAPI (blue). Scale bar = 50 pm. (D) Representative Western blot analysis showing Na^+^-K*-ATPase expression levels in HK-2 cells from the Normal, Hypoxia, and Hypoxia+3m-SMEF groups, β-actin was used as a loading control. *(E)* Densitometry analysis of Na*-K*-ATPase expression normalized on β-actin. Data are presented as mean ± SEM. N=2. Data were analyzed using one-way ANOVA. “ P < 0.01 was considered statistically significant.

### Application of the 3m-SMEF improves IRI-induced AKI in both male and female mice

Renal function was evaluated by measuring plasma creatinine levels 24 hours after IRI in both male and female mice. IRI resulted in a significant elevation in plasma creatinine (1.76 ± 0.08 mg/dL in males and 1.75 ± 0.09 mg/dL in females) compared to control (0.08 ± 0.01 mg/dL in males and 0.054 ± 0.007 mg/dL in females) and sham groups (0.154 ± 0.020 mg/dL in males and 0.126 ± 0.005 mg/dL in females). Application of 3m-SMEF dramatically reduced plasma creatinine by approximately 90% in males (0.23 ± 0.04 vs. 1.76 ± 0.08 mg/dL) and 95% in females (0.15 ± 0.02 vs. 1.75 ± 0.05 mg/dL), restoring values to near-sham levels) (Fig. 3A–B). These improvements substantially exceeded those previously achieved with the 3^rd^ generation SMEF (40% reduction in plasma creatinine). These results demonstrate that 3m-SMEF provides marked protection against IRI-induced functional decline in both sexes.

**Figure 3.**
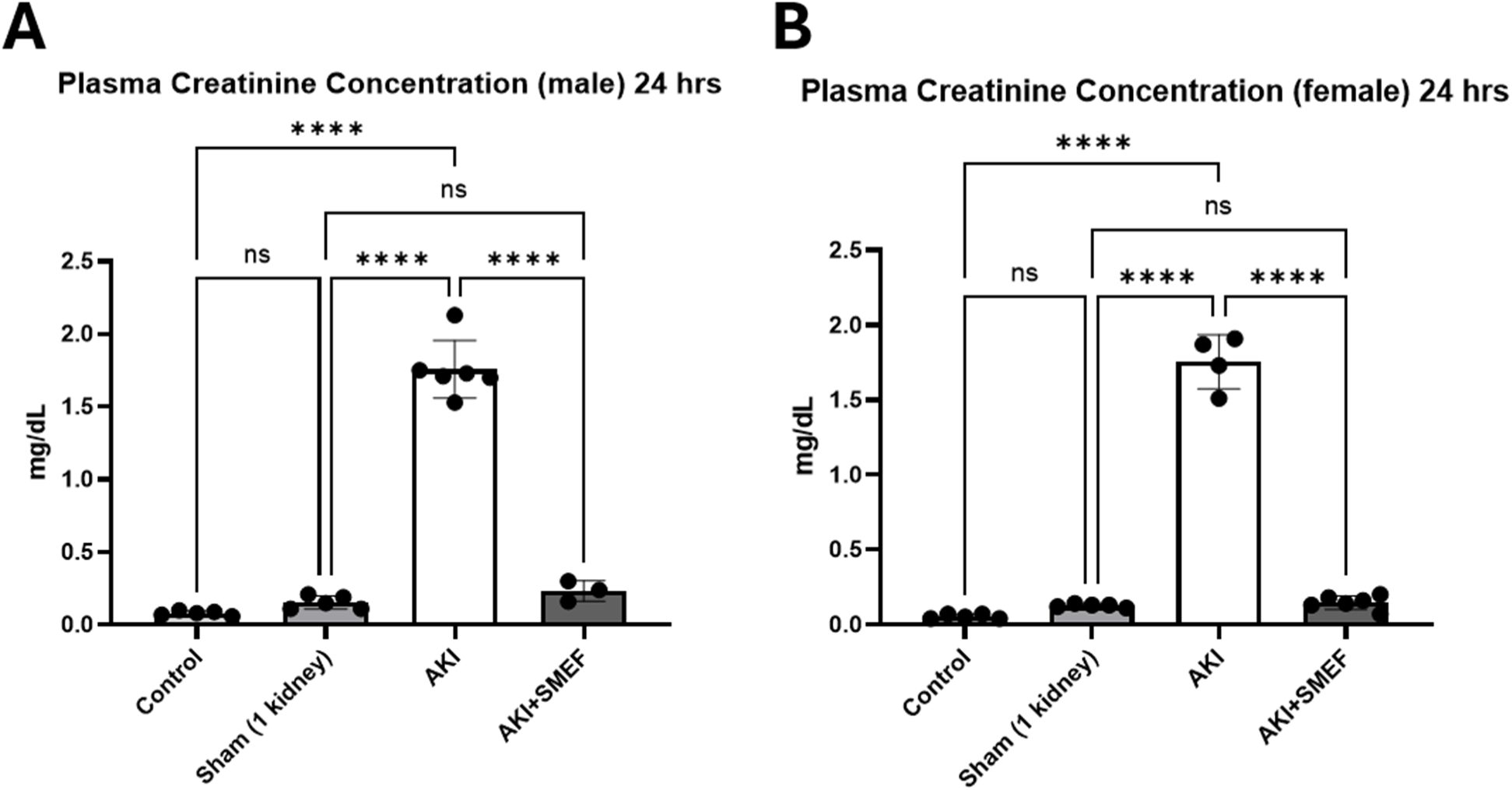
Plasma creatinine levels in male *(A)* and female *(B)* mice. Plasma creatinine concentrations were measured in C57BL/6J mice under the following experimental conditions: Control (no surgery), Sham (unilateral nephrectomy only), AKI (18 min renal ischemia), and AKI with 3m-SMEF applied to the kidney during 18 min of ischemia for mails and 24 min for females). Plasma samples were collected 24 h after ischemia-reperfusion injury (IRI). Data are presented as mean ± SEN. N=3-6. Data were analyzed using one-way ANOVA. **** P < 0.0001.

### Application of the 3m-SMEF restores GFR after IRI

GFR measured using transdermal FITC-sinistrin clearance was markedly reduced in both male (245.27 ± 19.00 vs. 7.33 ± 0.83 μL/min)and female AKI mice (250.11 ± 8.35 vs 27.79 ± 2.47 μL/min) compared with sham animals. Application of the 3m-SMEF significantly improved GFR in both male (208.00 ± 11.25 vs 159.48 ± 4.25 μL/min) and female AKI animals (217.90 ± 8.58 vs 159.48 ± 4.25 μL/min), with values approaching those of control and sham animals (Fig. 4A–B). These findings confirm that 3m-SMEF preserves renal filtration capacity following ischemic injury.

**Figure 4.**
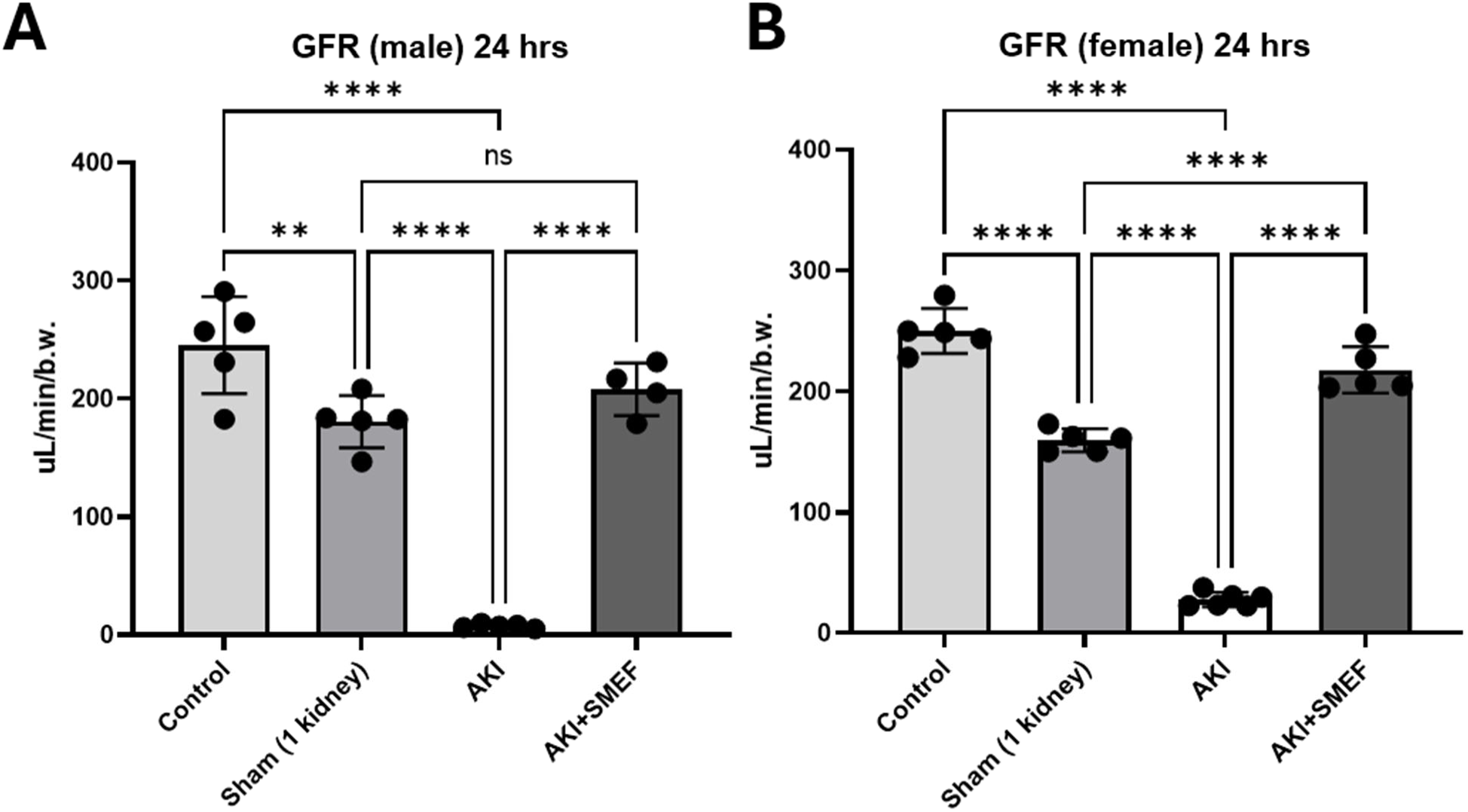
Glomerular filtration rate (GFR) measurements in male (A) and female (B) mice. Experimental groups included Control (no surgery), Sham (unilateral nephrectomy only), AKI (18 min renal ischemia), and AKI with 3m-SMEF applied to the kidney during the 18 min ischemic period (AKI+SMEF)). Data are presented as mean ± SEM. N=5-6. Data were analyzed using one-way ANOVA. ** P < 0.01, **** P < 0.0001.

### Application of the 3m-SMEF reduces kidney injury markers following IRI

Kidney injury molecule-1 (KIM-1) and neutrophil gelatinase-associated lipocalin (NGAL) levels were markedly increased in kidneys from AKI mice (271.24 ± 30.85 pg/mL and 10.73 ± 0.76 ng/mL, respectively) compared with those from normal control mice (60.14 ± 9.14 pg/mL and 0.59 ± 0.15 ng/mL, respectively). Treatment with 3m-SMEF significantly reduced both KIM-1 and NGAL levels (65.66 ± 18.52 pg/mL and 0.41 ± 0.12 ng/mL, respectively), restoring them to values comparable to those observed in normal control kidneys (Fig. 5A–B). These findings indicate that 3m-SMEF significantly attenuates IRI-induced tubular injury.

**Figure 5.**
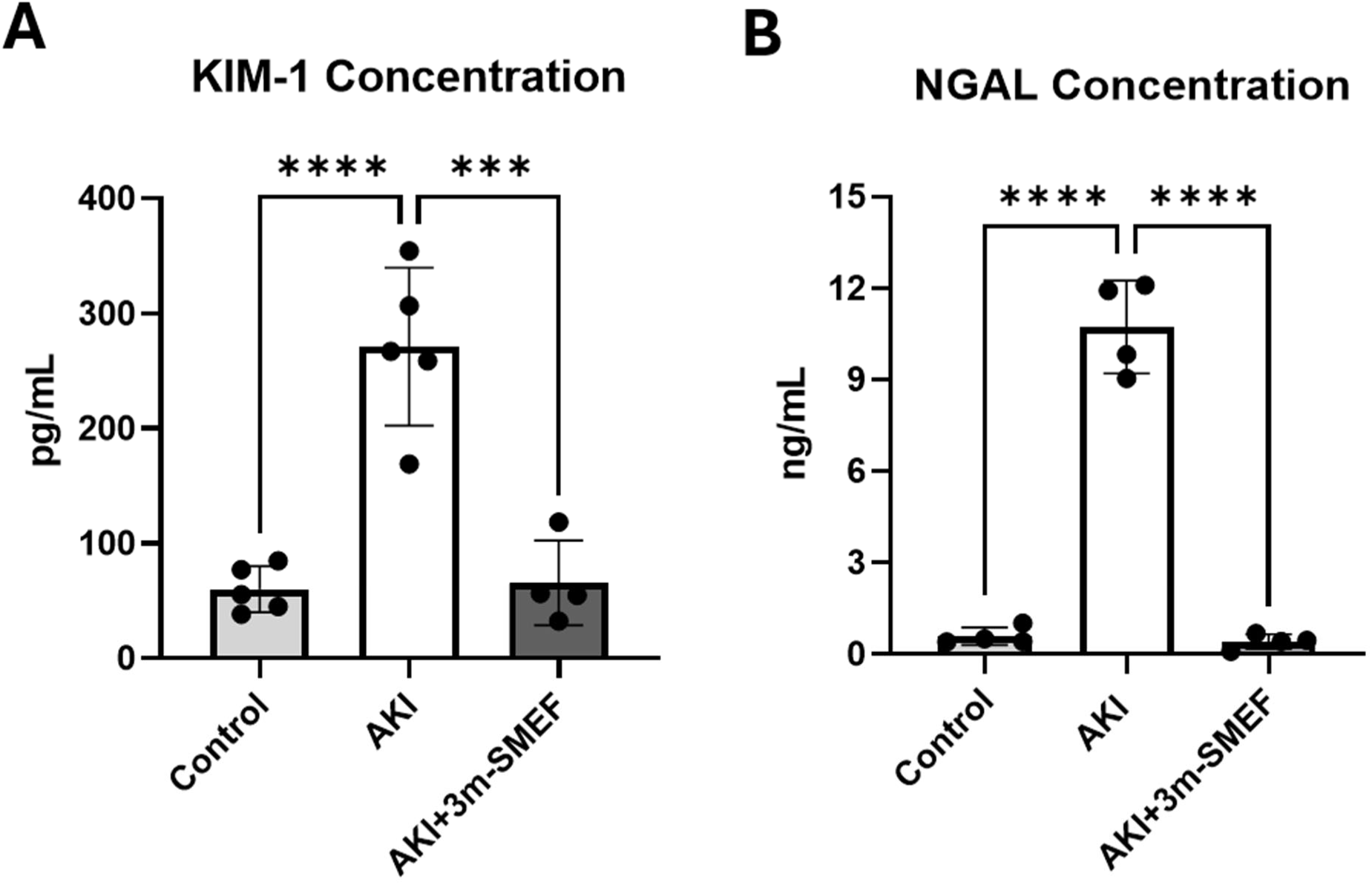
Kidney Injury Molecule-1 (KIM-1) and Neutrophil Gelatinase-Associated Lipocalin (NGAL) levels in male mice kidneys tissue. KIM-1 *(A)* and NGAL *(B)* levels were measured in sham kidney tissue (unilateral nephrectomy), AKI kidney tissue (mice subjected to 18 min of renal ischemia-reperfusion injury (AKI), and kidneys from mice with application of 3m-SMEF during the 18 min ischemic period (AKI+3m-SMEF). Kidney tissues were collected 24 h after injury, and protein levels were quantified using ELISA kits. Data are presented as mean ± SEM. N=5-6. Data were analyzed using one-way ANOVA. **** P < 0.0001

### Application of the 3m-SMEF preserves renal ATP levels following ischemic injury

Because the 3m-SMEF was designed to maintain the Na⁺/K⁺-ATPase activity independent of external ATP, the ATP levels should be preserved. Given that ATP depletion is a central feature of ischemic injury, we assessed renal tissue ATP levels to determine whether 3m-SMEF supported cellular energy preservation. IRI caused a pronounced reduction in ATP concentration (3.15 ± 0.28 ng/ml) compared to controls (17.18 ± 1.05 ng/ml). Application of the 3m-SMEF preserved ATP levels (13.40 ± 1.52 ng/ml), maintaining values close to those observed in the control group (Fig. 6). These results suggest that 3m-SMEF supports cellular energetics during ischemia and promotes the preservation of tubular cell function.

**Figure 6.**
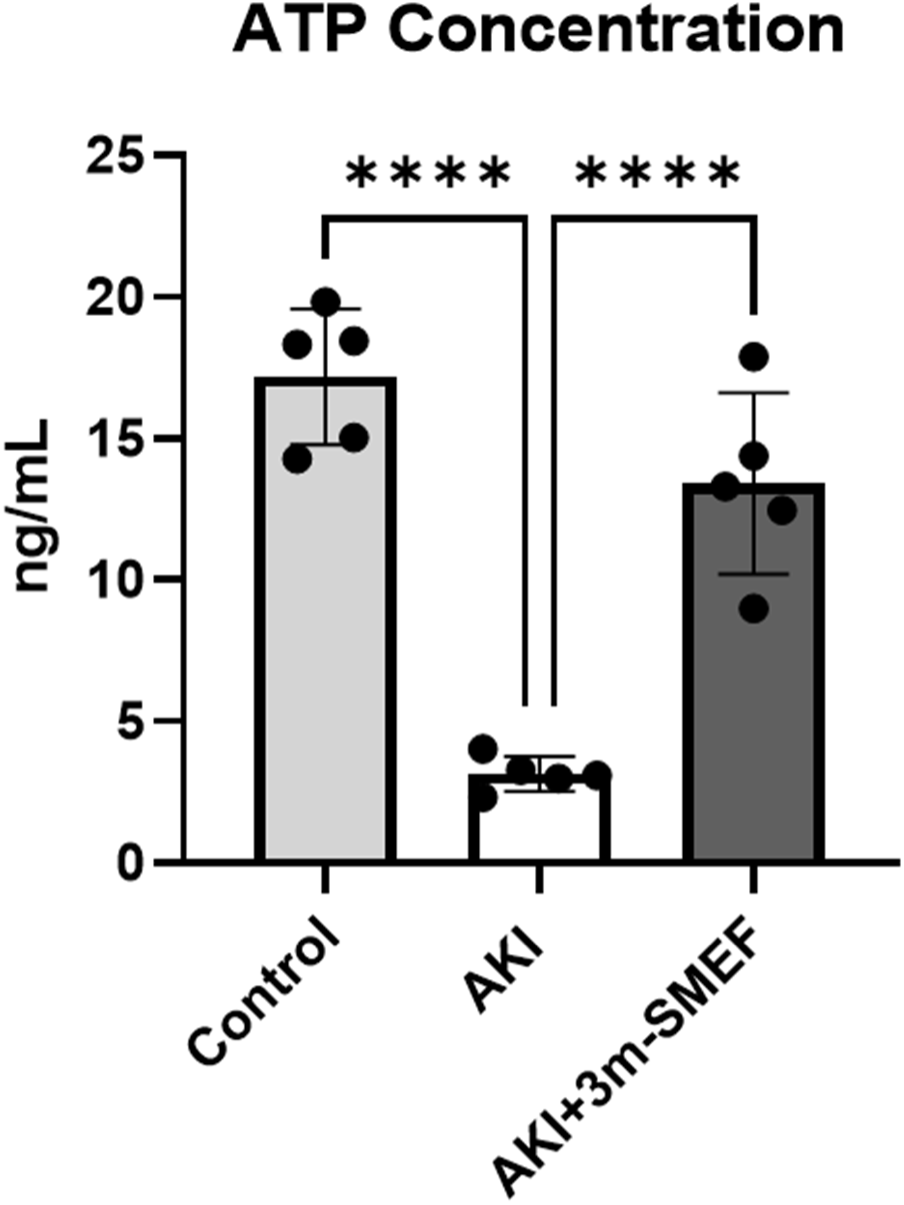
ATP levels in mouse kidney tissue. Control kidneys, AKI kidneys (mice subjected to renal ischemia-reperfusion injury) and kidneys from the mice where modified 3m-SMEF was applied during the ischemic period (AKI+3m-SMEF). Kidney tissues were collected 24 h after injury. Data are presented as mean ± SEM. N=5. Data were analyzed using one-way ANOVA. **** p < 0.0001.

### Application of the 3m-SMEF preserves Na⁺/K⁺-ATPase expression

To investigate the mechanism underlying 3m-SMEF-mediated protection, we examined Na⁺/K⁺-ATPase expression in kidney tissue. Western blot analysis revealed a reduction in α1 Na⁺/K⁺-ATPase expression in AKI mice compared to controls (>20-fold decrease) (Fig. 7 A, B). Application of the 3m-SMEF preserved Na⁺/K⁺-ATPase expression to levels comparable to the sham group (Fig. 7 A and B). These data suggest that 3m-SMEF prevented the degradation and translocation of the Na⁺/K⁺-ATPase, thereby stabilizing Na⁺/K⁺-ATPase expression during AKI.

**Figure 7.**
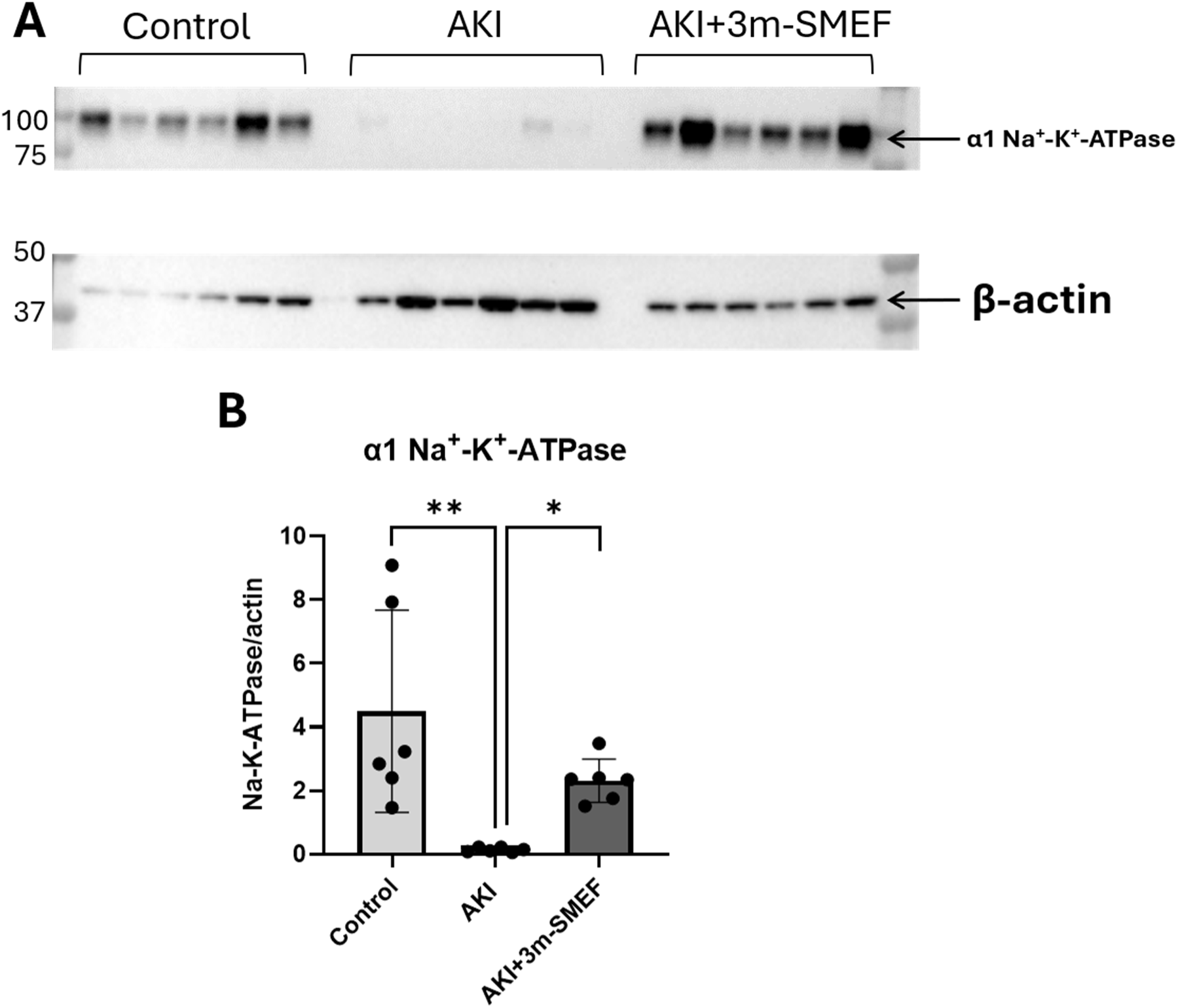
Na^+^/K^+^-ATPase protein expression in male mice kidneys. *(A)* Representative Western blot analysis of renal Na^+^/K^+^-ATPase α1 subunit with β-actin as the loading control in Control, AKI (18 min of ischemia), and AKI+3m-SMEF mice, in which 3m-SMEF was applied to the kidney during the 18-min ischemic period. *(B)* Band intensities were quantified by densitometry and normalized to β-actin. Data are presented as mean ± SEM. N=6. Statistical analysis was performed using an unpaired ttest with Welch’s correction. * P < 0.05, ** P < 0.01.

## Discussion

The present study demonstrates that the modified 3rd-generation Synchronization Modulation Electric Field (3m-SMEF) exhibits profound protective effects against both hypoxia-induced cell injury in vitro and IRI-induced AKI in vivo. In cultured HK-2 cells, 3m-SMEF reduced hypoxia-induced cytotoxicity and apoptosis while restoring Na⁺/K⁺-ATPase expression. In mice subjected to IRI, 3m-SMEF preserved renal function, as demonstrated by lowering of plasma creatinine, improvement in GFR, reduction in tubular injury markers KIM-1 and NGAL, restoration of renal ATP content, and maintenance of renal Na⁺/K⁺-ATPase expression at levels similar to normal control animals. These findings indicate that preservation of Na⁺/K⁺-ATPase-dependent cellular energy homeostasis is a fundamental mechanism of renal protection and that 3m-SMEF is a promising bioelectrical therapy for ischemic AKI.

The efficiency of the 3m-SMEF was first tested in HK-2 cells. The experiments in vitro provided direct evidence that 3m-SMEF protects renal tubular epithelial cells from hypoxic injury. Hypoxia markedly increased Cytotox Green and Annexin V staining, indicating enhanced cytotoxicity and activation of apoptosis, whereas 3m-SMEF significantly attenuated both responses. These findings are consistent with previous studies demonstrating that oxygen deprivation rapidly disrupts tubular epithelial cell homeostasis through ATP depletion, mitochondrial dysfunction, oxidative stress, and activation of regulated cell death pathways, ultimately leading to tubular injury and AKI progression ^12–19^. During renal ischemia, ATP depletion and mitochondrial dysfunction initiate a cascade of events that culminates in apoptosis, necrosis, and other forms of regulated cell death, accompanied by inflammatory activation and loss of tubular integrity ^17,18,20^. Therefore, the marked reduction in Cytotox Green and Annexin V staining observed in 3m-SMEF-treated HK-2 cells suggests that preservation of cellular bioenergetics by 3m-SMEF limits activation of multiple cell death pathways during hypoxic stress.

Another important finding was that 3m-SMEF restored Na⁺/K⁺-ATPase expression in hypoxic HK-2 cells. Remarkably, Na⁺/K⁺-ATPase expression exceeded that observed under normoxic conditions, suggesting that 3m-SMEF not only prevents hypoxia-induced loss of the pump but may also actively promote its recovery/stabilization and/or enhance its expression during cellular repair. Enhanced restoration of Na⁺/K⁺-ATPase could facilitate rapid re-establishment of transmembrane ion gradients, epithelial polarity, and vectorial sodium transport following hypoxic injury, thereby accelerating recovery of tubular function. Restoration of basolateral Na⁺/K⁺-ATPase localization is a critical determinant of functional recovery after ischemic injury ^21^. Although the underlying mechanisms remain unknown, this response may involve enhanced membrane trafficking, reduced protein degradation, or stabilization of existing Na⁺/K⁺-ATPase molecules ^22^. Further studies will be required to explore these possibilities.

Based on the promising in vitro findings, we explored whether they would be translate into preservation of whole-organ function in vivo. Renal IRI caused the expected deterioration of kidney function, as we previously reported ^10^, characterized by elevated plasma creatinine, reduced GFR, increased expression of KIM-1 and NGAL, depletion of renal ATP, and reduced renal Na⁺/K⁺-ATPase expression. Application of 3m-SMEF significantly attenuated each of these changes. These observations are consistent with previous studies demonstrating that ischemia-induced ATP depletion rapidly impairs Na⁺/K⁺-ATPase-dependent tubular transport, disrupts epithelial polarity, promotes mitochondrial dysfunction and inflammatory signaling, and ultimately reduces glomerular filtration and renal function ^2,23–26^. The restoration of ATP, together with the preservation of Na⁺/K⁺-ATPase, suggests that 3m-SMEF maintains tubular epithelial function during the critical early phase of reperfusion, thereby limiting the progression of AKI.

Na⁺/K⁺-ATPase is the major consumer of cellular ATP in renal tubular epithelial cells, where it is essential for maintaining transmembrane sodium gradients, membrane potential, epithelial polarity, cell volume, and vectorial transport ^27–29^. Meanwhile, Na⁺/K⁺-ATPase is the main driving force for secondary ion transport in the kidneys that controls salt and water excretion and maintains hemostasis of extracellular volume and electrolytes. Preservation of Na⁺/K⁺-ATPase by 3m-SMEF may maintain both cellular integrity and ion transport functions during ischemic injury. Consequently, loss of Na⁺/K⁺-ATPase during ischemia disrupts numerous downstream cellular processes beyond sodium transport. ATP depletion rapidly impairs Na⁺/K⁺-ATPase activity, resulting in intracellular sodium accumulation, potassium loss, calcium overload, mitochondrial dysfunction, oxidative stress, inflammatory activation, apoptosis, necrosis, and ultimately tubular epithelial cell death ^4,30^. Therefore, preservation of Na⁺/K⁺-ATPase represents an attractive therapeutic strategy for preventing ischemic kidney injury.

Our previous work demonstrated that synchronization-modulated electric fields maintained Na⁺/K⁺-ATPase activity under ATP-deficient conditions and reduced renal ischemic injury ^10^. The present study extends these observations by showing that the 3m-SMEF restores Na⁺/K⁺-ATPase protein expression in both cultured tubular cells and ischemic kidneys. Notably, Na⁺/K⁺-ATPase expression in 3m-SMEF-treated hypoxic cells exceeded basal levels, suggesting that 3m-SMEF actively promotes recovery of the transport machinery rather than simply preventing degradation. Thus, maintenance of Na⁺/K⁺-ATPase may represent a unique therapeutic mechanism underlying 3m-SMEF-mediated protection. The preservation of Na⁺/K⁺-ATPase likely explains many of the downstream protective effects observed in this study. Since Na⁺/K⁺-ATPase accounts for the majority of ATP consumption in renal tubular epithelial cells and drives active sodium reabsorption, preservation of pump activity supports intracellular ionic homeostasis and limits secondary mitochondrial injury ^28,31^. Improved mitochondrial function would be expected to decrease reactive oxygen species generation, preserve ATP production, and reduce activation of inflammatory signaling pathways. Consequently, tubular epithelial cells become less susceptible to apoptosis, necrosis, and necroptosis, leading to reduced release of damage-associated molecular patterns and diminished immune activation ^14,23,24,32,33^. The reduction in Annexin V and Cytotox staining observed in HK-2 cells, together with lower renal KIM-1 and NGAL expression in vivo, is consistent with this mechanism. Rather than acting on individual inflammatory or apoptotic pathways, 3m-SMEF appears to preserve an upstream regulator of cellular homeostasis. By maintaining Na⁺/K⁺-ATPase function during ischemia, 3m-SMEF may interrupt the cascade linking ATP depletion to mitochondrial dysfunction, oxidative stress, inflammation, and tubular cell death.

Despite advances in supportive care, there are currently no approved pharmacological therapies that directly prevent ischemic AKI. Numerous interventions targeting oxidative stress, inflammation, apoptosis, ferroptosis, or mitochondrial dysfunction have shown promising efficacy in experimental models but have failed to translate into effective clinical therapies, likely because they target downstream consequences of ischemia rather than the primary disturbance in cellular energy metabolism ^14,34–36^.

3m-SMEF represents a fundamentally different therapeutic strategy. Rather than inhibiting a single signaling pathway, 3m-SMEF preserves the energy-dependent Na⁺/K⁺-ATPase system that coordinates tubular transport and cellular homeostasis. Because electrical field therapy is non-pharmacological, it may avoid many of the systemic adverse effects associated with drug therapies and could potentially be applied prophylactically during planned ischemic events such as kidney transplantation, partial nephrectomy, cardiovascular surgery, or other procedures associated with transient renal ischemia. Future studies evaluating long-term renal outcomes, optimal treatment parameters, and safety in large-animal models will be essential before clinical translation.

Several limitations should be acknowledged. First, this study examined early injury after ischemia-reperfusion, and longer follow-up is needed to determine whether 3m-SMEF prevents progression from AKI to chronic kidney disease (CKD). Second, although Na⁺/K⁺-ATPase preservation was consistently observed, pump activity was not directly measured, and the molecular mechanisms underlying increased pump expression remain unclear. Future studies should determine whether 3m-SMEF regulates ATP1A1 transcription, protein trafficking, membrane localization, or protein stability. Third, mitochondrial function, oxidative stress, inflammatory signaling, and regulated cell death pathways were not directly measured and therefore remain mechanistic hypotheses requiring further investigation. In addition, it is not clear what the effect of 3m-SMEF is when it is applied during the reperfusion stage, which is more clinically relevant, rather than applied during the ischemic stage. Finally, although the mouse model closely reproduces ischemic AKI, validation in large-animal models (like swine models) and clinically relevant settings will be necessary before translation to patients.

Together, our findings identify preservation of Na⁺/K⁺-ATPase function and cellular bioenergetics as a novel therapeutic paradigm for ischemic AKI and provide compelling evidence that 3m-SMEF is a promising bioelectrical therapy for preventing renal ischemia-reperfusion injury.

## METHODS

### Modification of 3^rd^-generation SMEF (3m-SMEF)

To improve the efficacy of the third-generation Synchronization Modulation Electric Field platform, we modified the field application strategy, waveform, and parameter optimization. The new generation incorporates dual energy transformation functions and power injections to more efficiently generate ATP. The pumping rate and the synchronization of the pump molecules were also further optimized to minimize potential side effects on the cells. Because renal tubules are oriented in multiple directions, maximal electric-field effects on Na⁺/K⁺-ATPase occur only when the tubular membrane is perpendicular to the applied field. To improve whole-kidney coverage, four electrodes arranged as two orthogonal pairs were used to apply electric fields sequentially in the X- and Y-directions, alternating every three stimulation cycles. This approach increases the probability of synchronizing Na⁺/K⁺-ATPase molecules throughout the kidney. The protocol first synchronized Na⁺/K⁺-ATPase molecules at their intrinsic turnover frequency (50 Hz) with 10–20 stimulation pulses, then rapidly switched to a predetermined higher frequency to transiently accelerate pump activity. This synchronization-and-acceleration strategy promotes faster restoration of transmembrane ionic gradients and membrane potential during ischemia while reducing dependence on limited intracellular ATP, thereby improving preservation of renal function.

### HK-2 cell culturing, Hypoxia and Application of the 3m-SMEF to the cells

Human kidney proximal tubular epithelial cells (HK-2)(CRL-2190, ATCC) were cultured in complete growth medium DMEM/F-12 supplemented with 10% FBS, 100 U/mL penicillin, and 100 μg/mL streptomycin at 37°C in a humidified atmosphere containing 5% CO₂. At 70–80% confluence, cells were seeded onto 12-well Transwell inserts (0.4 μm polycarbonate membrane) and divided into three experimental groups. The 3m-SMEF (50–100 Hz, 100 mV) was applied using two silver electrodes positioned above and below the membrane, generating a uniform electric field across the cell monolayer in the apical-to-basal direction. Hypoxia was induced in HK2 cells in a modulator incubator chamber (Billups-Rothenberg, Del Mar, CA) by flushing with a 95% N2/5% CO2 gas mixture for 10 min and then sealing the chamber as described in the literature ^37,38^. Hypoxia was maintained for 24 hours, followed by a 2-hour reoxygenation (H/R) by opening the chamber and replacing the hypoxic medium with an oxygenated medium.

### Animals

Male and female C57BL/6J mice were used. The mice were, on average, 11 weeks of age and weighed 27 grams. They were procured from the Jackson Laboratory. Upon arrival, the animals were housed in a temperature-controlled environment with a 12-hour light/dark cycle, with free access to food and water. The animals were randomly assigned to control or treatment groups in accordance with the experimental requirements. All procedures and experiments were performed with the approval of the Institutional Animal Care and Use Committee at the University of South Florida College of Medicine and in accordance with all federal guidelines. Animals were euthanized in accordance with the policies of the American Veterinary Medical Association. Euthanasia was performed while the animals were under surgical anesthesia and was involved exsanguination by severing the heart and aorta. The death of the animals was confirmed before disposing of the carcasses.

### Renal IRI-AKI and Application of the 3m-SMEF to the kidney

Unilateral ischemia–reperfusion injury (IRI) was performed in male and female C57BL/6J mice as we previously described ^10^. In short, the mice were anesthetized with isoflurane (2–4%). A vertical midline abdominal incision was made to access the abdominal cavity. Following right nephrectomy, ischemia was induced by clamping the left renal pedicle with a microvascular clamp for 18 min in males and 20 min in female mice. Body temperature was maintained between 36.8°C and 37.2°C during surgery. In the treatment group, the 3m-SMEF was applied to the left kidney during the ischemic period using four electrodes arranged as two orthogonal pairs (X- and Y-directions) mounted on an elastic fabric that surrounded the kidney, ensuring uniform electric field coverage over the entire organ. At the end of the procedure, the SMEF electrodes were removed, the abdomen was closed, and the animals recovered.

### Plasma Creatinine

Plasma creatinine was measured at 24h post-surgery. The tail was snipped, and blood was collected from the tail caudal veins (40µL) with heparinized micro-hematocrit capillary tubes. The blood was transferred to EDTA-coated microcentrifuge tubes and centrifuged (10,000 g for 5 minutes at 4°C) to separate plasma (20 µL). Creatinine levels were measured at UAB O’Brien Center with LC-MS/MS.

### Glomerular filtration rate

GFR was measured in conscious mice at 24h post-surgery with FITC-sinistrin clearance rate using MediBeacon transdermal mini GFR monitoring devices, as previously described ^39^.

### Tissue Collection

At the experimental endpoint, mice designated for biochemical and molecular assays were euthanized. Kidneys were promptly removed, briefly rinsed in cold phosphate-buffered saline (PBS) to remove residual blood, and immediately flash-frozen in liquid nitrogen to preserve tissue integrity and enzymatic activity. Frozen samples were stored at −80°C until further processing and analysis.

### Kidney Injury Molecule-1 (KIM-1) and Neutrophil Gelatinase-Associated Lipocalin (NGAL)

Kidney Injury Molecule-1 (KIM-1) and Neutrophil Gelatinase-Associated Lipocalin (NGAL) levels were measured 24 hours post-ischemia-reperfusion injury (IRI) using commercially available ELISA kits: KIM-1 (Abcam, ab213477) and NGAL (Abcam, ab199083), following the manufacturers’ instructions as previously described ^10^. Kim-1 and NGAL concentrations were calculated from standard curves generated using the kit-provided standards and analyzed according to the respective manufacturer’s protocols.

### Measurement of ATP

Tissue ATP levels were measured 24 hours post-ischemia-reperfusion injury (IRI) using a commercially available ELISA kit according to the manufacturers’ instructions (Abcam, ab83355), as previously described ^10^. Briefly, kidney tissues were harvested, weighed, and immediately homogenized in the ATP assay buffer provided in the kit, followed by deproteinization and neutralization with the Deproteinizing Sample Preparation Kit (Abcam, ab204708) to eliminate enzymatic interference. The absorbance of the samples was taken at 570 nm using a microplate reader (Synergy HTX, BioTek). ATP concentrations (nmol/mg tissue) were calculated by interpolating sample absorbance from a standard curve generated with known ATP concentrations.

### Western blot analysis

Na⁺/K⁺-ATPase expression was analyzed by Western blot as previously described ^40^. Shortly, HK-2 cells and kidney tissues were lysed in RIPA buffer containing protease and phosphatase inhibitors. Protein concentrations were determined using the BCA assay. Equal amounts of protein (30 μg) were separated by 4–15% SDS-PAGE, transferred to PVDF membranes, blocked with 5% nonfat milk/TBST, and stained with primary antibodies against Na⁺/K⁺-ATPase (cs-2170) and then with secondary antibodies (GAR, Sigma, A6154). Equal protein loading was confirmed by β-actin (ab-6276) immunoblotting. Protein bands were visualized by enhanced chemiluminescence (ECL), and band intensities were quantified using Image Lab (Bio-Rad) or ImageJ programs.

### Statistical analysis

Statistical analysis was performed using Prism 10 (GraphPad Software). The data were compared using Unpaired t test with Welch’s correction or ordinary one-way ANOVA. Data are presented as means ± SE. * P < 0.05, ** P < 0.01, *** P < 0.001, **** P < 0.0001 were considered statistically significant.

## Author contributions

RL, NH, NZ, LW, WC designed experiments; NH, NZ, CE, TW, NHS, IL, WC performed experiments and analyzed the data; NH and NZ prepared the original draft; NH, NZ, CE, TW, CP, IL, NHS, LW, WC and RL reviewed and edited the manuscript, RL supervised the study. All authors reviewed, revised, and approved the article.

## Data availability

All data generated or analyzed during this study are included in this published article.

## References

1 Doyle, J. F. & Forni, L. G. Acute kidney injury: short-term and long-term effects. Crit Care 20, 188 (2016). 10.1186/s13054-016-1353-y

2 Bonventre, J. V. & Yang, L. Cellular pathophysiology of ischemic acute kidney injury. J Clin Invest 121, 4210–4221 (2011). 10.1172/JCI45161

3 Kellum, J. A. et al. Acute kidney injury. Nat Rev Dis Primers 7, 52 (2021). 10.1038/s41572-021-00284-z

4 Molitoris, B. A. Na(+)-K(+)-ATPase that redistributes to apical membrane during ATP depletion remains functional. Am J Physiol 265, F693–697 (1993). 10.1152/ajprenal.1993.265.5.F693

5 Venkatachalam, M. A., Weinberg, J. M., Kriz, W. & Bidani, A. K. Failed Tubule Recovery, AKI-CKD Transition, and Kidney Disease Progression. J Am Soc Nephrol 26, 1765–1776 (2015). 10.1681/ASN.2015010006

6 Han, W. K., Bailly, V., Abichandani, R., Thadhani, R. & Bonventre, J. V. Kidney Injury Molecule-1 (KIM-1): a novel biomarker for human renal proximal tubule injury. Kidney Int 62, 237–244 (2002). 10.1046/j.1523-1755.2002.00433.x

7 Mishra, J. et al. Neutrophil gelatinase-associated lipocalin (NGAL) as a biomarker for acute renal injury after cardiac surgery. Lancet 365, 1231–1238 (2005). 10.1016/S0140-6736(05)74811-X

8 Levin, M. Molecular bioelectricity: how endogenous voltage potentials control cell behavior and instruct pattern regulation in vivo. Mol Biol Cell 25, 3835–3850 (2014). 10.1091/mbc.E13-12-0708

9 Funk, R. H., Monsees, T. & Ozkucur, N. Electromagnetic effects - From cell biology to medicine. Prog Histochem Cytochem 43, 177–264 (2009). 10.1016/j.proghi.2008.07.001

10 Chen, W. et al. Reducing ischemic kidney injury through application of a synchronization modulation electric field to maintain Na(+)/K(+)-ATPase unctions. Sci Transl Med 14, eabj4906 (2022). 10.1126/scitranslmed.abj4906

11 Semenza, G. L. Hypoxia-inducible factors in physiology and medicine. Cell 148, 399–408 (2012). 10.1016/j.cell.2012.01.021

12 Wheaton, W. W. & Chandel, N. S. Hypoxia. 2. Hypoxia regulates cellular metabolism. Am J Physiol Cell Physiol 300, C385–393 (2011). 10.1152/ajpcell.00485.2010

13 Flood, D., Lee, E. S. & Taylor, C. T. Intracellular energy production and distribution in hypoxia. J Biol Chem 299, 105103 (2023). 10.1016/j.jbc.2023.105103

14 Li, H. et al. Advances in understanding the role of mitochondria in renal ischemia-reperfusion injury. Clin Exp Nephrol 29, 1685–1698 (2025). 10.1007/s10157-025-02727-3

15 Pan, Y. & Zhu, J. Mitochondria in Renal Ischemia-Reperfusion Injury: From Mechanisms to Therapeutics. Biomedicines 14 (2026). 10.3390/biomedicines14020310

16 Chen, Y. et al. Mitochondrial metabolism and targeted treatment strategies in ischemic-induced acute kidney injury. Cell Death Discov 10, 69 (2024). 10.1038/s41420-024-01843-5

17 Noh, M. R. & Padanilam, B. J. Cell death induced by acute renal injury: a perspective on the contributions of accidental and programmed cell death. Am J Physiol Renal Physiol 327, F4–F20 (2024). 10.1152/ajprenal.00275.2023

18 Han, S. J. & Lee, H. T. Mechanisms and therapeutic targets o ischemic acute kidney injury. Kidney Res Clin Pract 38, 427–440 (2019). 10.23876/j.krcp.19.062

19 Pefanis, A., Ierino, F. L., Murphy, J. M. & Cowan, P. J. Regulated necrosis in kidney ischemia-reperfusion injury. Kidney Int 96, 291–301 (2019). 10.1016/j.kint.2019.02.009

20 Sanz, A. B., Sanchez-Nino, M. D., Ramos, A. M. & Ortiz, A. Regulated cell death pathways in kidney disease. Nat Rev Nephrol 19, 281–299 (2023). 10.1038/s41581-023-00694-0

21 Stoops, E. H. & Caplan, M. J. Trafficking to the apical and basolateral membranes in polarized epithelial cells. J Am Soc Nephrol 25, 1375–1386 (2014). 10.1681/ASN.2013080883

22 Dada, L. A. et al. Phosphorylation and ubiquitination are necessary for Na,K-ATPase endocytosis during hypoxia. Cell Signal 19, 1893–1898 (2007). 10.1016/j.cellsig.2007.04.013

23 Malek, M. & Nematbakhsh, M. Renal ischemia/reperfusion injury; from pathophysiology to treatment. J Renal Inj Prev 4, 20–27 (2015). 10.12861/jrip.2015.06

24 Sancho-Martinez, S. M., Lopez-Novoa, J. M. & Lopez-Hernandez, F. J. Pathophysiological role of different tubular epithelial cell death modes in acute kidney injury. Clin Kidney J 8, 548–559 (2015). 10.1093/ckj/sfv069

25 Schrezenmeier, E. V., Barasch, J., Budde, K., Westhoff, T. & Schmidt-Ott, K. M. Biomarkers in acute kidney injury - pathophysiological basis and clinical performance. Acta Physiol (Oxf) 219, 554–572 (2017). 10.1111/apha.12764

26 Zuk, A. & Bonventre, J. V. Acute Kidney Injury. Annu Rev Med 67, 293–307 (2016). 10.1146/annurev-med-050214-013407

27 Katz, A. I. Renal Na-K-ATPase: its role in tubular sodium and potassium transport. Am J Physiol 242, F207–219 (1982). 10.1152/ajprenal.1982.242.3.F207

28 Feraille, E. & Doucet, A. Sodium-potassium-adenosinetriphosphatase-dependent sodium transport in the kidney: hormonal control. Physiol Rev 81, 345–418 (2001). 10.1152/physrev.2001.81.1.345

29 Rajasekaran, S. A., Barwe, S. P. & Rajasekaran, A. K. Multiple functions of Na,K-ATPase in epithelial cells. Semin Nephrol 25, 328–334 (2005). 10.1016/j.semnephrol.2005.03.008

30 Dong, Y. et al. Ischemic Duration and Frequency Determines AKI-to-CKD Progression Monitored by Dynamic Changes of Tubular Biomarkers in IRI Mice. Front Physiol 10, 153 (2019). 10.3389/fphys.2019.00153

31 Mandel, L. J. Primary active sodium transport, oxygen consumption, and ATP: coupling and regulation. Kidney Int 29, 3–9 (1986). 10.1038/ki.1986.2

32 Zhao, Z. et al. XJB-5-131 inhibited ferroptosis in tubular epithelial cells after ischemia-reperfusion injury. Cell Death Dis 11, 629 (2020). 10.1038/s41419-020-02871-6

33 Fan, H., Liu, J., Sun, J., Feng, G. & Li, J. Advances in the study of B cells in renal ischemia-reperfusion injury. Front Immunol 14, 1216094 (2023). 10.3389/fimmu.2023.1216094

34 Szeto, H. H. Pharmacologic Approaches to Improve Mitochondrial Function in AKI and CKD. J Am Soc Nephrol 28, 2856–2865 (2017). 10.1681/ASN.2017030247

35 Feng, Q. et al. Ferroptosis and Acute Kidney Injury (AKI): Molecular Mechanisms and Therapeutic Potentials. Front Pharmacol 13, 858676 (2022). 10.3389/fphar.2022.858676

36 Li, Z. et al. The Pathogenesis of Ischemia-Reperfusion Induced Acute Kidney Injury Depends on Renal Neutrophil Recruitment Whereas Sepsis-Induced AKI Does Not. Front Immunol 13, 843782 (2022). 10.3389/fimmu.2022.843782

37 Malhotra, R., Tyson, D. W., Rosevear, H. M. & Brosius, F. C., 3rd. Hypoxia-inducible factor-1alpha is a critical mediator of hypoxia induced apoptosis in cardiac H9c2 and kidney epithelial HK-2 cells. BMC Cardiovasc Disord 8, 9 (2008). 10.1186/1471-2261-8-9

38 Martinive, P. et al. Impact of cyclic hypoxia on HIF-1alpha regulation in endothelial cells--new insights for anti-tumor treatments. FEBS J 276, 509–518 (2009). 10.1111/j.1742-4658.2008.06798.x

39 Li, M. et al. Macula densa-specific NOS1 knockout determines susceptibility to ischemic acute kidney injury. Clin Sci (Lond) 140, 589–605 (2026). 10.1042/CS20258807

40 Zheleznova, N. N., Kumar, V., Kurth, T. & Cowley, A. W., Jr. Hydrogen peroxide (H(2)O(2)) mediated activation of mTORC2 increases intracellular Na(+) concentration in the renal medullary thick ascending limb of Henle. Sci Rep 11, 7300 (2021). 10.1038/s41598-021-86678-1

